## Supplementary figures and images for "Inflammatory response in hematopoietic stem and progenitor cells triggered by activating SHP2 mutations evokes blood defects"

### Fig 1-supplement 1

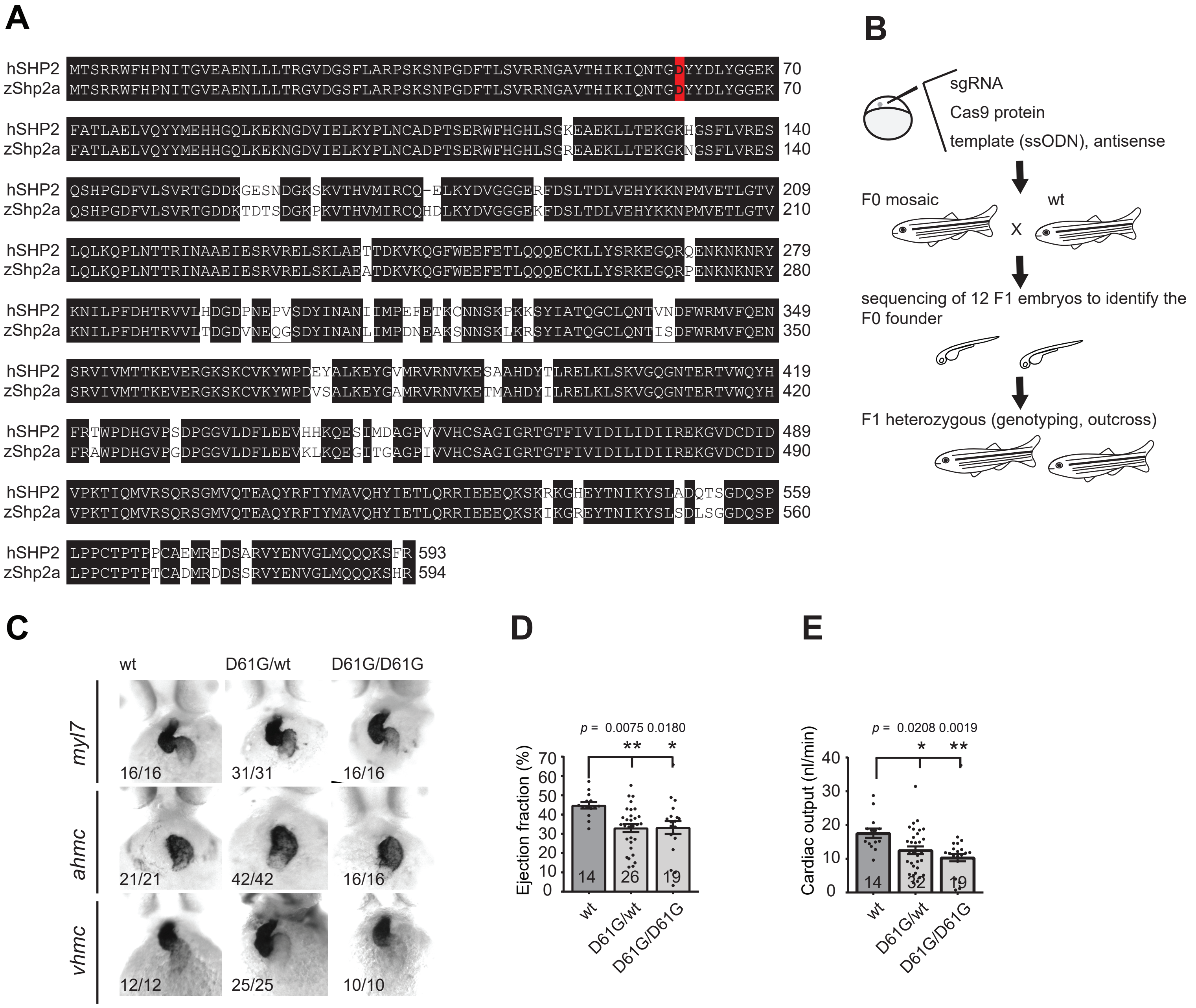

### Fig 2-supplement 1

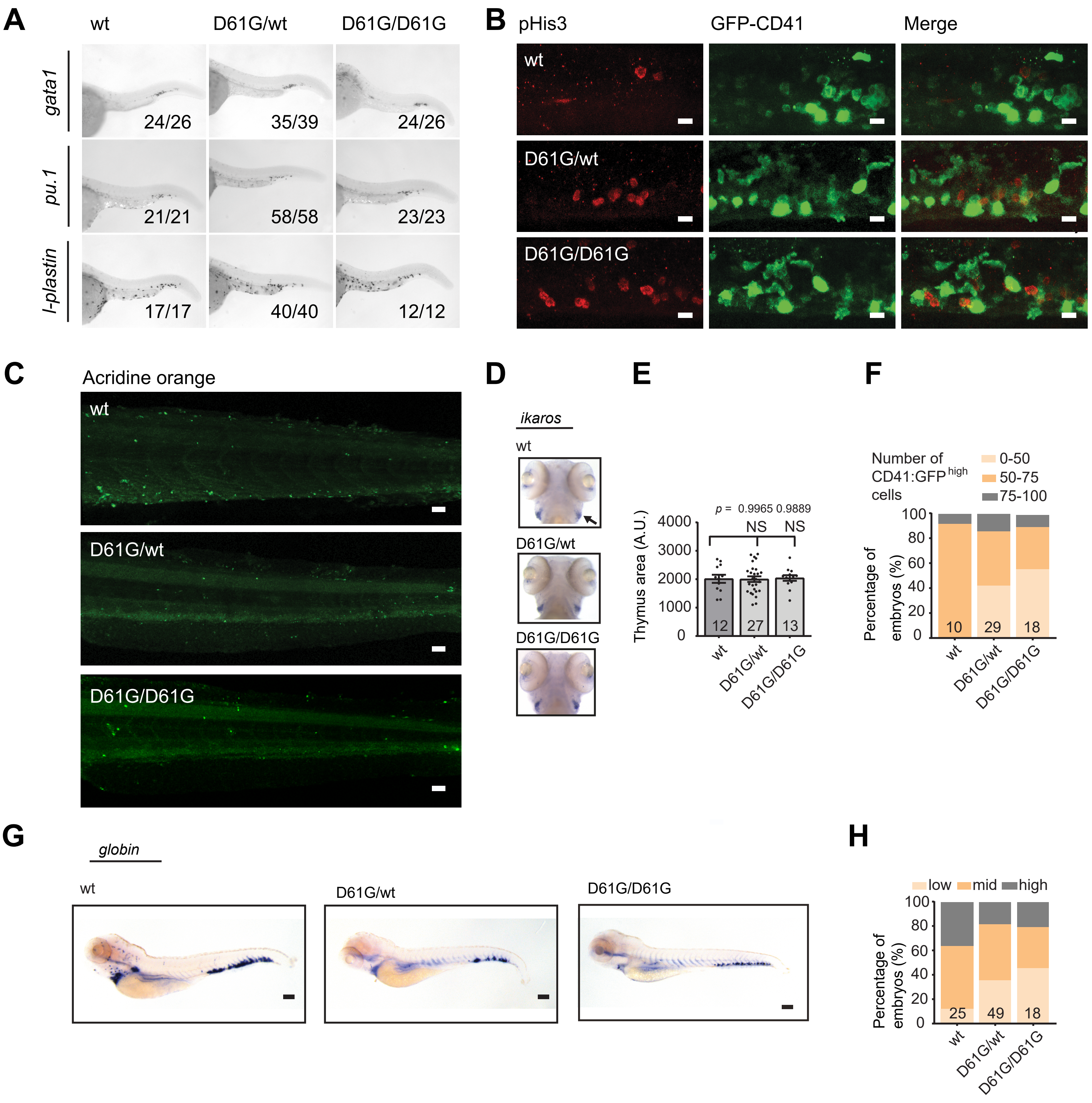

### Fig 3-supplement 1

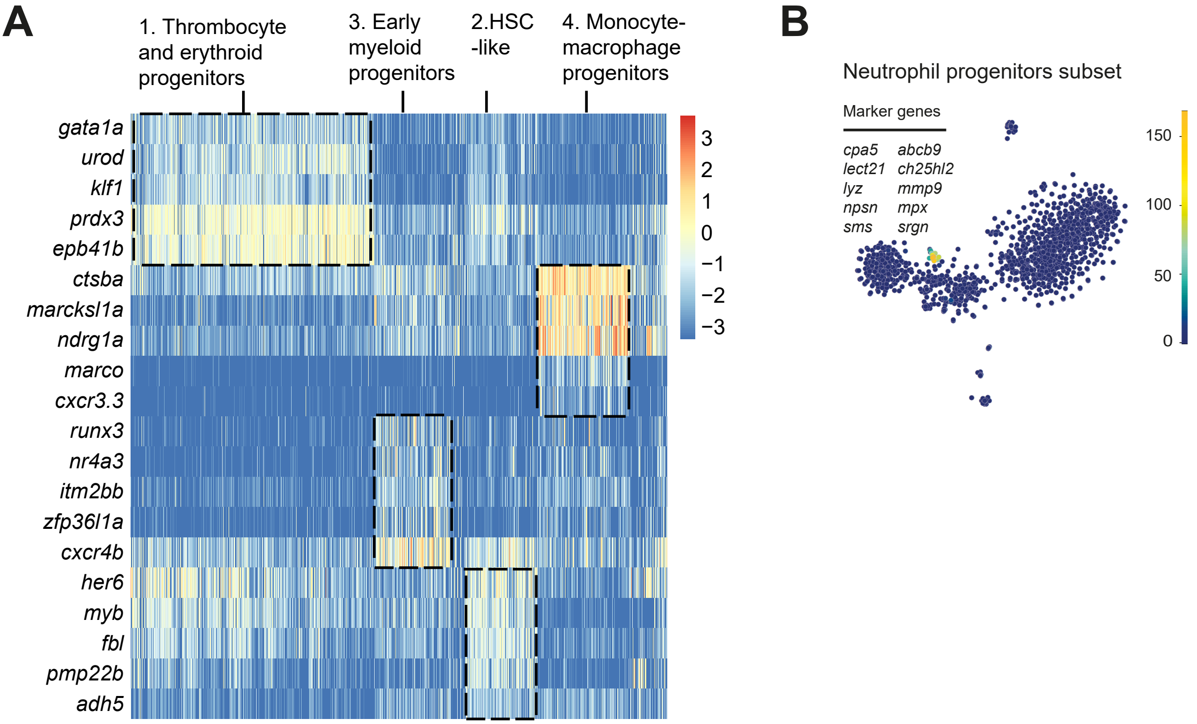

### Fig 4-supplement 1

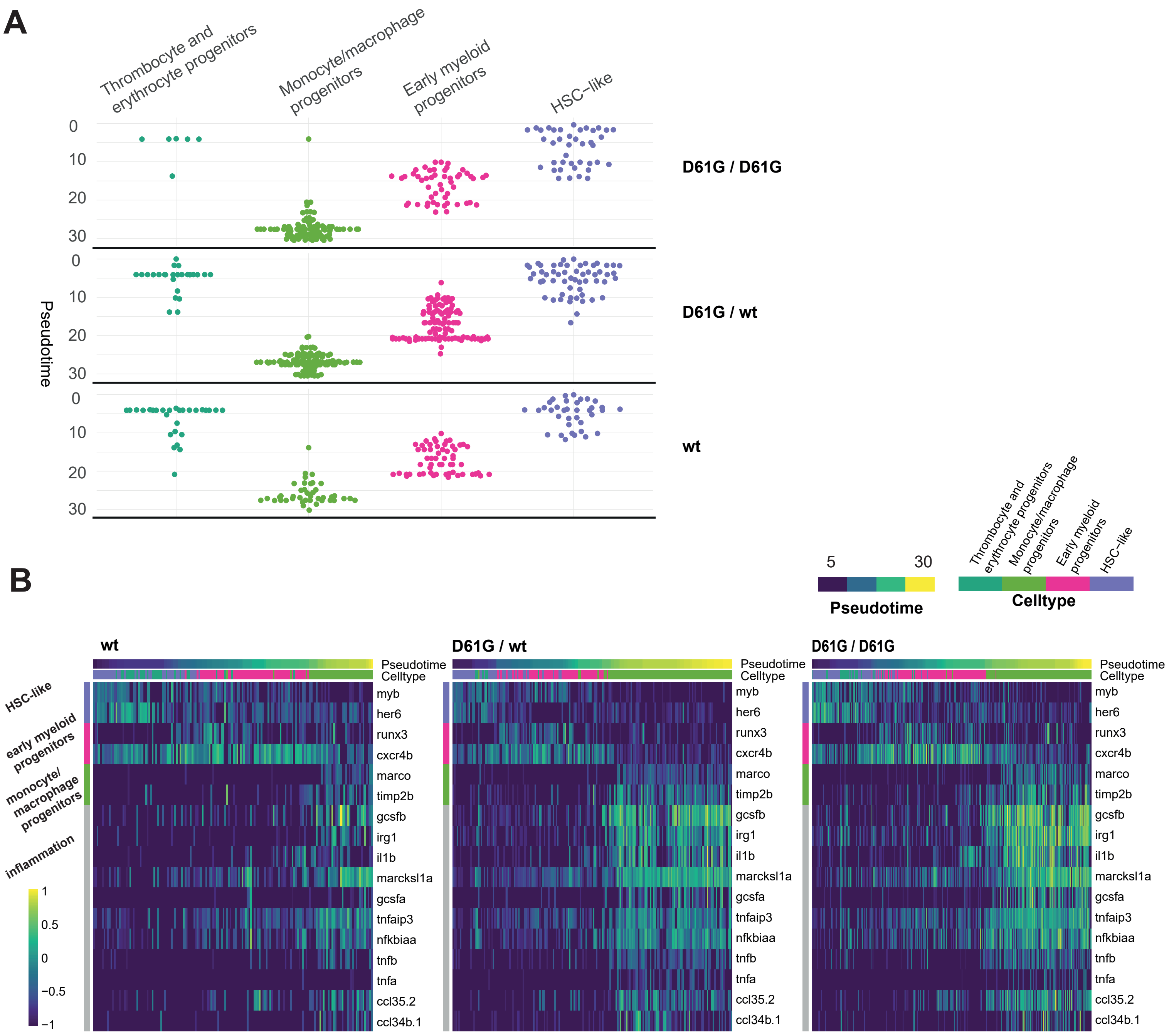
