## Supplementary figure legends for "Inflammatory response in hematopoietic stem and progenitor cells triggered by activating SHP2 mutations evokes blood defects"

**Figure 1-figure supplement 1. Defective heart function, but not heart morphology in Shp2$^{D61G}$ zebrafish embryos. (A)** Sequence alignment of human SHP2 and zebrafish Shp2a polypeptides performed by ClustalW. The residue D61 is highlighted in red, while the identical amino acids are highlighted in black. The amino acid number is marked to the right of each row. **B)** Schematic representation of the procedure used to establish the Shp2$^{D61G}$ mutant line, as described in the Materials and methods section. **(C)** Representative images of the WISH staining for *myl7*, *ahmc* and *vhmc* expression in the hearts of 3dpf old Shp2$^{wt}$, Shp2$^{D61G/wt}$ and Shp2$^{D61G/D61G}$ zebrafish embryos. Numbers in the pictures indicate the number of embryos with the phenotype represented in the image. Ejection fraction **(D)** and cardiac output **(E)** determined from the high speed videos of the heart of 5dpf old Shp2$^{wt}$ , Shp2$^{D61G/wt}$ and Shp2$^{D61G/D61G}$ zebrafish embryos. Measurements originate from at least three distinct experiments. Numbers on the bars depict the number of embryos. Error bars represent standard error of the mean (SEM).*p < 0.05; **p < 0.01, ***p < 0.001, NS, non significant, ANOVA complemented by Tukey HSD.

**Figure 2-figure supplement 1. Hematopoiesis in Shp2D61G zebrafish. (A)** Representative images of the WISH staining for gata1, pu.1 and l-plastin expression in the tail region of 48hpf old Shp2$^{wt}$ , Shp2$^{D61G/wt}$ and Shp2$^{D61G/D61G}$ zebrafish embryos. Scale bar is 250µm. Numbers in the pictures indicate the number of embryos with the phenotype represented in the image. **(B)** Cell proliferation was assessed in the CHT region of 5 dpf old Shp2$^{wt}$ , Shp2$^{D61G/wt}$ and Shp2$^{D61G/D61G}$ embryos in the Tg(cd41:GFP) background by immunohistochemistry using antibodies specific for pHis3 and GFP. Cells positive for both GFP and pHis3 are indicated with white arrows in the Merge panel. Scale bar is 10µm. **(C)** Representative images of the Acridine orange staining of the CHT region of the 5dpf old Shp2$^{wt}$ , Shp2$^{D61G/wt}$ and Shp2$^{D61G/D61G}$ embryos. Scale bar is 50µm. **(D)** WISH of 5dpf Shp2$^{wt}$ , Shp2$^{D61G/wt}$ and Shp2$^{D61G/D61G}$ embryos using ikaros specific probe. Thymus is indicated (arrow). **(E)** Size of ikaros-positive thymus, NS, non significant, ANOVA complemented by Tukey HSD. **(F)** Number of cd41:GFP$^{high}$ cells in Shp2$^{wt}$ , Shp2$^{D61G/wt}$ and Shp2$^{D61G/D61G}$ zebrafish embryos at 5dpf were counted and percentage of embryos with either 0-50, 50-75 or 75-100 cd41:GFP$^{high}$ cells was plotted. **(G)** WISH of 5dpf Shp2$^{wt}$ , Shp2$^{D61G/wt}$ and Shp2$^{D61G/D61G}$ embryos using β-globin specific probe. Scale bar,150µm. **(H)** Quantification of β-globin expression in Shp2$^{wt}$ , Shp2$^{D61G/wt}$ and Shp2$^{D61G/D61G}$ embryos scored as low, mid and high. **(E,F,H)** Measurements originate from at least three distinct experiments. Numbers on the bars depict the number of embryos.

**Figure 3-figure supplement 1. Identification of different HSPCs subpopulations based on differential expression of representative genes. (A)** Heat map showing scaled expression [log TPM (transcripts per million) values] of representative genes (y-axis) in all cells (x-axis). Cells belonging to distinct clusters are squared with a dashed line and cluster identity is indicated on the top of the heat map. Cluster 1 is characterized by expression of genes characteristic for thrombocyte and erythroid differentiation, such as *gata1a* and *klf1*. Cells from cluster 2 express typical hematopoietic stem cell markers, such as *c-myb* and *her6* (*HES1* analogue in human). Cells from cluster 3 express early myeloid progenitor markers, such as *runx3* and *cxcr4b*. Cells from cluster 4 express genes typical for both monocyte and macrophages, such as *marco* and *ctsba*. **(B)** tSNE maps showing the sum of total read-counts of selected neutrophil specific genes.

**Figure 4-figure supplement 1. Distinct expression profiles of mutant zebrafish embryos in pseudotime (A)** Identities of the four clusters of HSPCs of Shp2$^{wt}$ , Shp2$^{D61G/wt}$ and Shp2$^{D61G/D61G}$ genotypes from the trajectory in Fig. 4, ordered in pseudotime. **(B)** Heat map showing scaled expression [log TPM (transcripts per million) values] of representative genes (y-axis) in cells of Shp2$^{wt}$ , Shp2$^{D61G/wt}$ and Shp2$^{D61G/D61G}$ genotypes from the selected trajectory route, ordered along the pseudotime (x-axis). The cell type and pseudotime are annotated above the heat map.
