## Supplementary Table 2 for "Inflammatory response in hematopoietic stem and progenitor cells triggered by activating SHP2 mutations evokes blood defects"

### Characteristics of JMML PTPN11 mutated patients and healty subjects submitted in RNAseq.

Transcripts : PTPN11(NM_002834.3), ASXL1(NM_015338.5)

| JMML Patients | JMML_153-04 | JMML_131-07 | JMML_166-15 | JMML_178-20 | JMML_190-24 | | |
|---|---|---|---|---|---|---|---|
| Gender | M | M | M | M | M | | |
| Age at diagnosis (years) | 0,4 | 3 | 4 | 0,3 | 4 | | |
| Platelets ($10^9$/L) at diagnosis | 112 | 64 | 40 | 122 | 62 | | |
| WBC ($10^9$/L) at diagnosis | 37 | 13 | 16 | 51 | 35 | | |
| Monocytes ($10^9$/L) at diagnosis | 7,7 | 1,2 | 1,5 | 6,9 | 6,6 | | |
| HbF elevated for age | No | Yes | Yes | No | Yes | | |
| *LIN28B* overexpression | No | Yes | Yes | Yes | Yes | | |
| Pulmonary involvement | No | No | No | No | Yes | | |
| HSCT | No | Yes | Yes | Yes | Yes | | |
| Blast crisis | No | No | No | No | No | | |
| Relapse post HSCT | No | No | No | No | No | | |
| Death | No | No | No | No | No | | |
| Variant of *PTPN11* | p.Ala72Thr | p.Gly503Val | p.Glu76Gln | p.Asp61Tyr | p.Glu76Lys | | |
| VAF of initiating mutation | 45% | 42% | 42% | 44% | 46% | | |
| Additional mutations | | | *PTPN11*: p.Thr468Met | | *ASXL1*: p.Gly646Trpfs*12 | | |
| VAF of initiating mutation | | | 42% | | 32% | | |
| Sorted fractions submitted to RNA sequencing | HSC, CMP, GMP, MEP | CMP, GMP, MEP | HSC, MPP, CMP, GMP, MEP | MPP, CMP, GMP, MEP | CMP, GMP, MEP | | |
| **Healthy subjects** | **BM_04** | **BM_05** | **BM_06** | **BM_09** | **BM_23** | **BM_24** | **BM_27** |
| Gender | M | F | F | M | F | F | F |
| Age (years) | 2,2 | 2,7 | 9,4 | 6,8 | 8,5 | 5 | 7,2 |
| Sorted fractions submitted to RNA sequencing | HSC, CMP, GMP, MEP | HSC, CMP, GMP, MEP | HSC, CMP, GMP, MEP | GMP, MEP | HSC | HSC | MPP |
